# Stool-derived eukaryotic RNA biomarkers for detection of high-risk adenomas

**DOI:** 10.1101/534412

**Authors:** Erica Barnell, Yiming Kang, Andrew Barnell, Katie Campbell, Kimberly R. Kruse, Elizabeth M. Wurtzler, Malachi Griffith, Aadel A. Chaudhuri, Obi L. Griffith

## Abstract

**Background and aims:** Colorectal cancer (CRC) is the second leading cause of cancer related deaths in the United States. Mortality is largely attributable to low patient compliance with screening and a subsequent high frequency of late-stage diagnoses. Noninvasive methods, such as stool- or blood-based diagnostics could improve patient compliance, however, existing techniques cannot adequately detect high-risk adenomas (HRAs) and early-stage CRC.

**Methods:** Here we apply cancer profiling using amplicon sequencing of stool-derived eukaryotic RNA for 275 patients undergoing prospective CRC screening. A training set of 154 samples was used to build a random forest model that included 4 feature types (differentially expressed amplicons, total RNA expression, demographic information, and fecal immunochemical test results). An independent hold out test set of 121 patients was used to assess model performance.

**Results:** When applied to the 121-patient hold out test set, the model attained a receiver operating characteristic (ROC) area under the curve (AUC) of 0.94 for CRC and a ROC AUC of 0.87 for CRC and HRAs. In aggregate, the model achieved a 91 % sensitivity for CRC and a 73% sensitivity for HRAs at an 89% specificity for all other findings (medium-risk adenomas, low-risk adenomas, benign polyps, and no findings on a colonoscopy).

**Conclusion:** Collectively, these results indicate that in addition to early CRC detection, stool-derived biomarkers can accurately and noninvasively identify HRAs, which could be harnessed to prevent CRC development for asymptomatic, average-risk patients.

## Introduction

Colorectal cancer (CRC) is the third most common cancer in both men and women in the United States, the second deadliest cancer globally, and accounted for over 50,000 deaths in 2018 in the US alone.^1,2^ Disease onset is typically insidious, starting as a small polyp which can take several years to further accrue somatic mutations and develop into an invasive carcinoma.^3,4^ If detected early, CRC has a five-year survival rate of 92%. However, 63% of newly diagnosed patients have advanced disease, with an associated five-year survival rate as low as 14%.^1,5,6^ Late-stage diagnosis typically results from patient noncompliance with screening guidelines, indicating these cancers could have been detected earlier by following standard protocols. CRC screening compliance has remained stagnant over the past 20 years and is currently estimated to be approximately 60%,^1,7^ well below the National Colorectal Cancer Roundtable’s goal of 80%.^7^ Additionally, nearly a quarter of adults in at-risk populations have never been screened.^6^ Historically, low compliance rates have been due to the inconvenience, unpleasantness, and perceived hazards of colonoscopies. In an open-ended survey of 660 patients regarding the most important barrier to CRC screening, three of the top four responses cited were directly related to the invasive nature of a colonoscopy: “afraid/fear,” “prep unpleasant,” and/or “anticipated pain”.^8^ Therefore, it is likely that a noninvasive alternative screening method would significantly increase the number of patients opting for CRC screening.

While many noninvasive tests have been developed to address compliance issues, none compare to the diagnostic accuracy of a colonoscopy.^9^ As such, colonoscopies remain the preferred first-line screening option by gastroenterologists. Currently, the most accurate noninvasive diagnostic on the market (Exact Sciences, Cologuard) cites a CRC sensitivity of 92%.^10^ However, the high-risk adenoma (HRA) detection rate for this diagnostic is only 42%.^10^ Other noninvasive stool-based tests include the fecal occult blood test and the fecal immunochemical test (FIT), which use lateral flow for detection of blood in stool. These alternatives can be highly sensitive (79%) and specific (94%) for CRC, but have HRA sensitivity of less than 30%.^11^ Accurate detection of precancerous adenomas would allow for preemptive excision of dysplastic tissue prior to carcinogenesis, thus reducing the CRC incidence and associated morbidity and mortality.^4,12^

Given the wide range of genomic variants that can cause healthy tissue to transform into a premalignant lesion, traditional diagnostics that target a small number of recurrent variants show reduced sensitivity for early carcinoma and precancerous change.^4^ The broad genomic landscape for CRC lesions can potentially be captured by evaluating a panel of RNA biomarkers that encompass the universal effects of most precancerous variants.^13,14^ Evaluating the downstream molecular symptoms derived from precancerous variants could improve sensitivity for adenomas.^15–17^ However, analysis of human RNA biomarkers in stool samples is extremely challenging due to extensive RNA degradation and a high bacterial transcript burden.^18,19^ Previous studies report that 25%-50% of stool samples evaluated for eukaryotic RNA are inadequate for downstream analysis.^20^

We describe herein a method to reliably extract and evaluate stool-derived eukaryotic RNA (seRNA) transcripts. We first use these biomarkers to identify transcripts associated with CRC and HRAs in a prospective screening population. Subsequently, we build an algorithm that incorporates differentially expressed amplicons, total RNA expression, demographic information, and results from a FIT to determine an individual’s risk for colonic lesions (CRC and HRA). We then validate the algorithm using an independent hold out test set, thus demonstrating proof-of-concept capability to noninvasively, sensitively, and specifically detect CRC and HRAs in a screening population (**Figure 1**).

**Figure 1.**
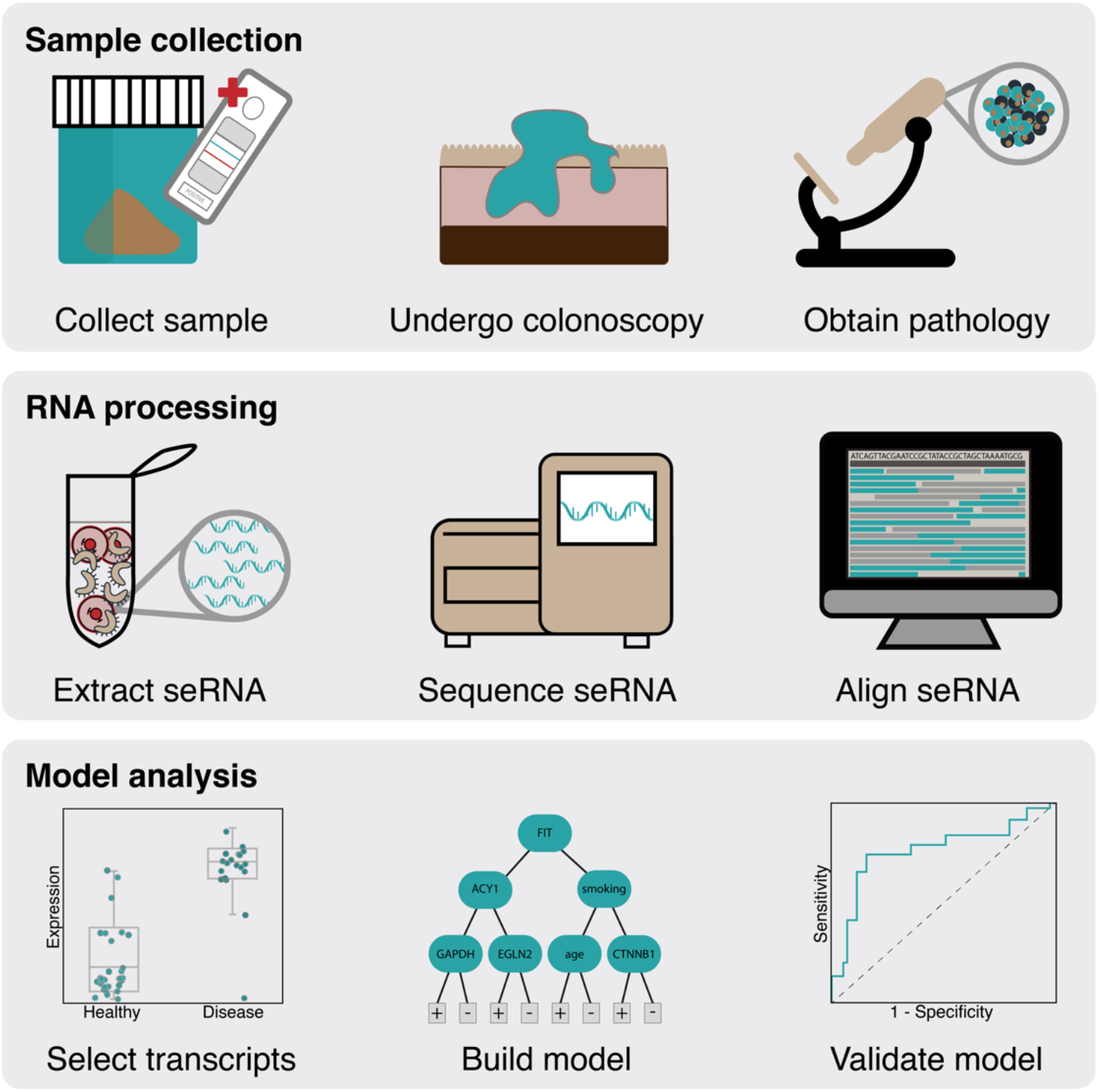
Overview of methods for obtaining samples, isolating stool-derived eukaryotic RNA (seRNA), sequencing seRNA, building a machine learning model, and testing the model’s accuracy. <u>Sample collection:</u> Stool samples were collected from patients prior to undergoing screening colonoscopy and subsequent histopathology. <u>RNA processing:</u> Eukaryotic transcripts were isolated from stool samples, expression analysis was performed via amplicon sequencing, and raw transcripts were aligned to the reference genome for quantification. <u>Model analysis:</u> Differentially expressed transcripts were identified, a random forest model was built, and the model was evaluated using an independent hold out test set. *Abbreviations: FIT–fecal immunochemical test; seRNA – stool-derived eukaryotic RNA*.

## Methods

### Study design

This study required prospective collection of stool samples from patients undergoing CRC screening or retrospective collection of stool samples from patients diagnosed with stage I-IV CRC prior to treatment or surgical resection. The Washington University School of Medicine (WUSM) (St. Louis, MO) Institutional Review Board (IRB) approved protocol and research procedures (IRB #20111107). The primary outcome of the study was to determine the feasibility of using stool-derived eukaryotic RNA (seRNA) to assess risk for CRC and HRAs. FITs were obtained for all samples prior to seRNA extraction. Subsequently, isolated seRNA was subjected to targeted amplification (TruSeq Targeted RNA Custom Panel; Illumina, San Diego, CA) and next-generation sequencing (NextSeq 550; Illumina, San Diego, CA). Sequencing reads were used for transcript selection, model development, and model validation (**see Supplementary Methods**).

### Transcript selection for model development

Feature selection was performed using bootstrapping of the training set (n = 154 samples). Specifically, the training set was segregated into 100 different 9:1 splits whereby each split was assessed for informative amplicons. An amplicon was considered informative if the absolute log2 fold-change was greater than 1 in both contrast groups (HRA vs. LRAs, benign polyps, no findings on a colonoscopy; MRAs vs. LRAs, benign polyps, no findings on a colonoscopy) and the ANOVA between the contrast groups had a p-value <0.05. If an amplicon was deemed informative in at least 33% of all bootstrapped splits, it was considered differentially expressed and eligible as a feature for model development. Other features (e.g., demographic information and raw GAPDH values) were also eligible for model development.

### Machine learning model development and assessment for high-risk adenomas

A random forest model was built using all 154 samples in the training set and all eligible features. Specifically, after feature selection, 5,000 decision trees were constructed using an independent bootstrapping procedure on the training samples; each node split was optimized by Gini Importance; each tree was built until it reached full depth. Output from the model provided a prediction between 0-1 whereby a larger number reflected increased confidence in a positive finding. Receiver operating characteristic (ROC) curves were created using model predictions and area under the curve (AUC) was used to measure model performance. For ROC curves plotted with the FIT, a positive FIT forced model prediction to equal to 1. The random forest model was employed on the 110 prospectively collected stool samples (excluding 11 retrospectively collected CRC samples). ROC curves were plotted with and without incorporating FIT results.

### Downsampling analysis of training set

To understand the extent of model training, downsampled fractions of training data were selected and performance was assessed using the testing set. The downsampling fractions ranged from 30% to 100% with 10% increments. For each downsampling fraction, feature selection was performed using bootstrapping, a random forest model was trained using the differentially expressed features, and the model was employed on the hold out test set. The resulting ROC AUC was used to assess model performance. This process was repeated 10 times for each downsampling fraction to reduce selection bias in subsampling, and model performance was assessed with and without incorporating FIT results.

### Ultimate model performance on the hold out test set and extrapolation to screening population

The random forest model was ultimately employed on all 121 patients in the complete hold out test set (110 prospectively collected samples and 11 retrospectively collected CRC samples). Output from the model provided a prediction between 0-1 and a positive FIT forced model prediction to equal 1. A ROC curve was plotted whereby only CRC samples were considered positive and all other categories were considered negative. A separate ROC curve was plotted whereby CRC and HRA samples were considered positive and all other categories were considered negative.

To attain a better approximation of ultimate model performance, the accuracy profile observed on the hold out test set was extrapolated to the relative frequencies expected in a prospective screening population. ROC curves as described above were plotted to show model performance. A point on the ROC curve was selected to optimize sensitivity and specificity. Subsequently, the blended sensitivity for CRC and HRAs, negative predictive value (NPV), and positive predictive value (PPV) were calculated.

## Results

### Sample demographic information

In total, stool samples from 275 individuals were collected for this study. Sequencing data, demographic information (i.e., gender, age, ethnicity, smoking status, and family history of CRC), results from a FIT, and colonoscopy results with histopathology information, if applicable, were obtained for all patients. In the study 11 patients had CRC (stage I-IV), 26 patients had high-risk adenomas (HRAs), 37 patients had medium-risk adenomas (MRAs), 61 patients had low-risk adenomas (LRAs), 50 patients had benign polyps, and 90 patients had no findings on a colonoscopy (**Table 1**). A training set of 154 patients was used for feature selection and model development and a hold out test set of 121-patients was used for model validation. There were two statistically significant differences between the characteristics of the training set and the hold out test set. First, retrospectively collected samples (i.e., samples from patients with CRC) were not included in the training set. Second, the hold out test set had different processing quality relative to the training set. Specifically, there was a reduction in the average stool input used for seRNA extraction (12.9 grams vs. 12.0 grams; p-value = 0.03), there was a reduction in the average seRNA concentration (168.6 ng/uL vs. 56.1 ng/uL; p-value < 0.01), and there was a reduction in average library preparation fragment size (200.6 base pairs vs. 192.2 base pairs; p-value < 0.01) (**Table 1**).

**Table 1.** Patient types, demographics and processing metrics associated with the prospective training set, the prospective hold out test set, the retrospective hold out test set, and the whole study cohort

| Category | Description | Prospective training set | Prospective hold out test set | Retrospective hold out test set | Total |
| --- | --- | --- | --- | --- | --- |
| CRC (1.0) | Stage I-IV CRC | 0 | 0 | 11 | 11 |
| HRA (2.1) | Any adenoma with carcinoma in situ or high-grade dysplasia | 1 | 2 | 0 | 3 |
| HRA (2.2) | Adenoma, villous growth pattern ($\geq 25\%$), any size | 4 | 2 | 0 | 6 |
| HRA (2.3) | Adenoma, $\geq 1.0$ cm in size | 5 | 5 | 0 | 10 |
| HRA (2.4) | Serrated lesion, $\geq 1.0$ cm in size | 5 | 2 | 0 | 7 |
| MRA (3) | $< 3$ adenoma(s), between 5-10 mm in size, non-advanced | 11 | 9 | 0 | 20 |
| MRA (4) | $\geq 3$ adenomas, $< 1.0$ cm in size, non-advanced | 10 | 7 | 0 | 17 |
| LRA (5) | 1 or 2 adenoma(s), $\leq 5$ mm in size, non-advanced | 37 | 24 | 0 | 61 |
| Benign polyps (6.1) | Hyperplastic, benign polyp(s) | 26 | 24 | 0 | 50 |
| No findings (6.2) | No findings on a colonoscopy | 55 | 35 | 0 | 90 |
| **Total** |  | **154** | **110** | **11** | **275** |

| Category | Description | Training set | Hold out test set | p-value* |
| --- | --- | --- | --- | --- |
| **Demographics** | Gender (Female vs. male) | 63.0% | 62.0% | 0.87 |
| | Age | $52.8 \pm 6.4$ | $54.3 \pm 7.4$ | 0.07 |
|  | Ethnic background (white vs. other) | 54.5% | 44.6% | 0.10 |
|  | Smoking status (never smoker vs. other) | 70.1% | 66.9% | 0.57 |
|  | Family history (negative family history vs. other) | 70.1% | 76.0% | 0.28 |
| **Processing metrics** | Average stool input (grams) | $12.9 \pm 3.2$ | $12.0 \pm 3.4$ | 0.03 |
| | Average RNA Integrity Value (RIN) | $4.6 \pm 1.2$ | $4.4 \pm 1.6$ | 0.18 |
| | Average RNA concentration - Qubit (ng/uL) | $168.6 \pm 139.8$ | $56.1 \pm 79.3$ | $< 0.01$ |
| | Average fragment size (bps) | $200.6 \pm 9.6$ | $192.2 \pm 16.8$ | $< 0.01$ |
| | Total library quantity (ng) | $29.7 \pm 29.5$ | $32.8 \pm 64.1$ | 0.60 |

\*significant change in frequency was calculated using a two-tailed chi square test
\*significant change in population mean was calculated using a two-tailed t-test

### Custom amplicon panel development

A custom panel of 639 amplicons was developed for target enrichment in the Illumina DesignStudio (**Figure 2A**). The custom amplicons were associated with 408 transcripts, which were selected using previously conducted research (**see Supplementary Methods**).^21^ Specifically, using microarray data (Affymetrix Human Transcriptome Array 2.0) from 265 individuals (177 patients with CRC or adenomas and 88 patients with no findings on a colonoscopy), 214 transcripts were identified as differentially expressed (p<0.03). Using molecular barcoding data (NanoString nCounter) from 85 individuals (59 with CRC or adenomas and 26 with no findings on a colonoscopy), an additional 123 transcripts were identified as differentially expressed. Finally, 71 transcripts were selected for the custom amplicon panel based on literature review.^22–26^

**Figure 2.**
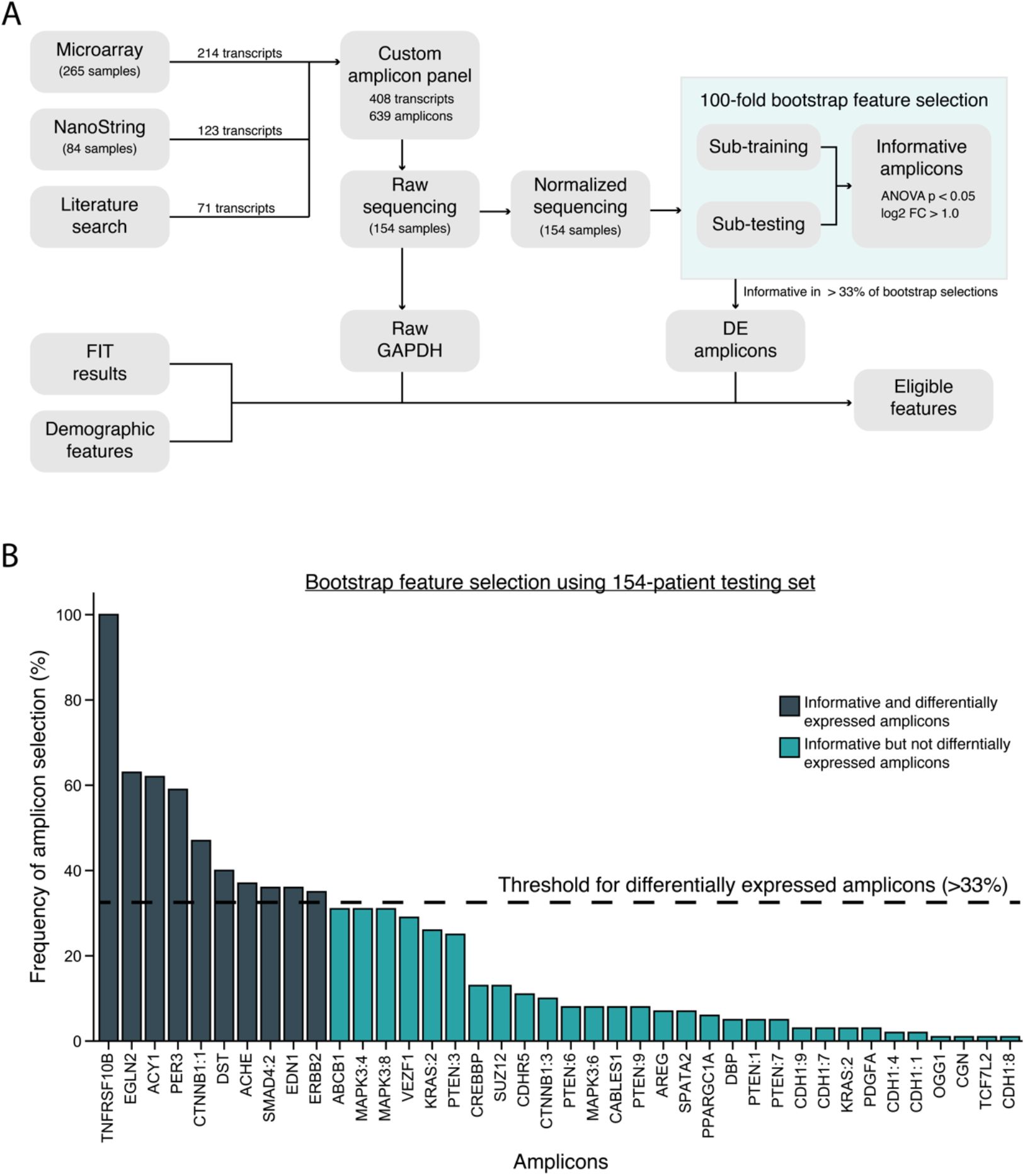
Eligible feature selection using bootstrapping of the training set. Transcripts used in the custom amplicon panel were selected based on a 265-sample microarray study, an 84-sample NanoString study, and pertinent literature. The 639 amplicons in the custom panel were used to obtain raw sequencing reads for the 154 samples in the training set. Amplicon expression was normalized to GAPDH and differentially expressed amplicons were identified using a bootstrapping method. Specifically, the 154-patient training set was split 100 times (9:1 sub-training vs. sub-testing) and each sub-training set was used to determine informative amplicons. An amplicon was considered informative if the absolute log2 fold-change between contrast groups was greater than 1 and the ANOVA p-value was less than 0.05. If an amplicon was observed in at least 33% of all 100 splits, then it was considered differentially expressed (DE) and was eligible as a feature for model development. Eligible features included results from a fecal immunochemical test (FIT), demographic features, raw GAPDH expression, and DE amplicons. **B)** If an amplicon was observed in at least 33% of all 100 splits (bootstrap threshold), then it was considered differentially expressed and was eligible as a feature for the final model. In total, 10 amplicons were selected as differentially expressed. *Abbreviations: FC – Fold change; ANOVA – analysis of variance; FIT – Fecal immunochemical test; DE – Differentially expressed*

### Technical and biological replicate analysis

Five patients were used to assess reproducibility of seRNA extraction, library preparation, and sequencing (**see Supplementary Methods**). Technical replicates consisting of aliquots from the same library preparation exhibited minimal difference in amplicon expression (Pearson r^2^ correlation range = 0.98-1.00; Pearson r^2^ average = 0.99) (**Supplementary Figure 1**). Replicates consisting of aliquots derived from a single stool sample had higher variation but exhibited good overall correlation. For the five biological replicates that underwent paralleled extraction, were subjected to different targeted enrichment strategies (200 ng with 30 cycles vs. 400 ng with 28 cycles), and were sequenced together, the average Pearson r^2^ correlation for expression of all 639 amplicons was 0.76 (**Supplementary Figure 2**). For the five biological replicates that underwent parallelled extraction, were subjected to the same target enrichment strategy (400 ng with 28 cycles), and were sequenced separately, the average Pearson r^2^ correlation for expression of all 639 amplicons was 0.73 (**Supplemental Figure 3**).

### Significant transcripts associated with disease

Normalized expression of 639 amplicons was evaluated for all samples in the training set (n = 154 samples). Of these 639 amplicons, 48 amplicons were not expressed in any sample and an additional 71 amplicons were not expressed in >95% of samples; these amplicons were eliminated from the analysis. For the remaining amplicons, a bootstrap analysis was performed (**see Methods**). In total, there were 40 amplicons identified as informative in at least 1 of the 100 splits and there were 10 amplicons identified as differentially expressed (informative in at least 33 of the 100 splits) (**Figure 2B**). For the 10 differentially expressed amplicons, the linkage to tumorigenesis and precancerous transformation are described in **Supplementary Table 1**. Ultimately, the 10 differentially expressed amplicons, raw *GAPDH* values, and 2 demographic identifiers (age and smoking status) were used as features for model development (**Figure 2A**).

### Accuracy profile for HRAs using 110-patient hold out test set (excluding 11 CRC patients)

The primary goal of this study was to use seRNA data to identify HRAs with a high sensitivity. Model performance was assessed through accuracy of predictions on the prospective hold out test set (excluding 11 retrospectively collected CRC stool samples). Using the 13 features described in **Supplementary Table 1**, a random forest model was built using all 154 samples in the training set and employed on the prospective hold out test set (**Figure 3A**). The most influential features in the final model were *ACY1* and *TNFRSF10B* (Gini Importance ≥ 0.13) and the least important feature was *PER3* (Gini Importance < 0.05). Raw *GAPDH* was the 4th most important feature in building the random forest model (**Supplementary Table 1**). The model attained a ROC AUC of 0.67 without FIT results and a ROC AUC of 0.78 when including FIT results (**Figure 3B**). It was observed that model output was correlated with disease severity, which was not provided as a feature for model training (**Figure 3C**). Specifically, directionality of categories (e.g., medium-risk adenomas are less severe than high-risk adenomas) and disease subsets (e.g., high-risk adenoma stratification based on adenoma size or level of dysplasia) were not available to the model during development.

**Figure 3.**
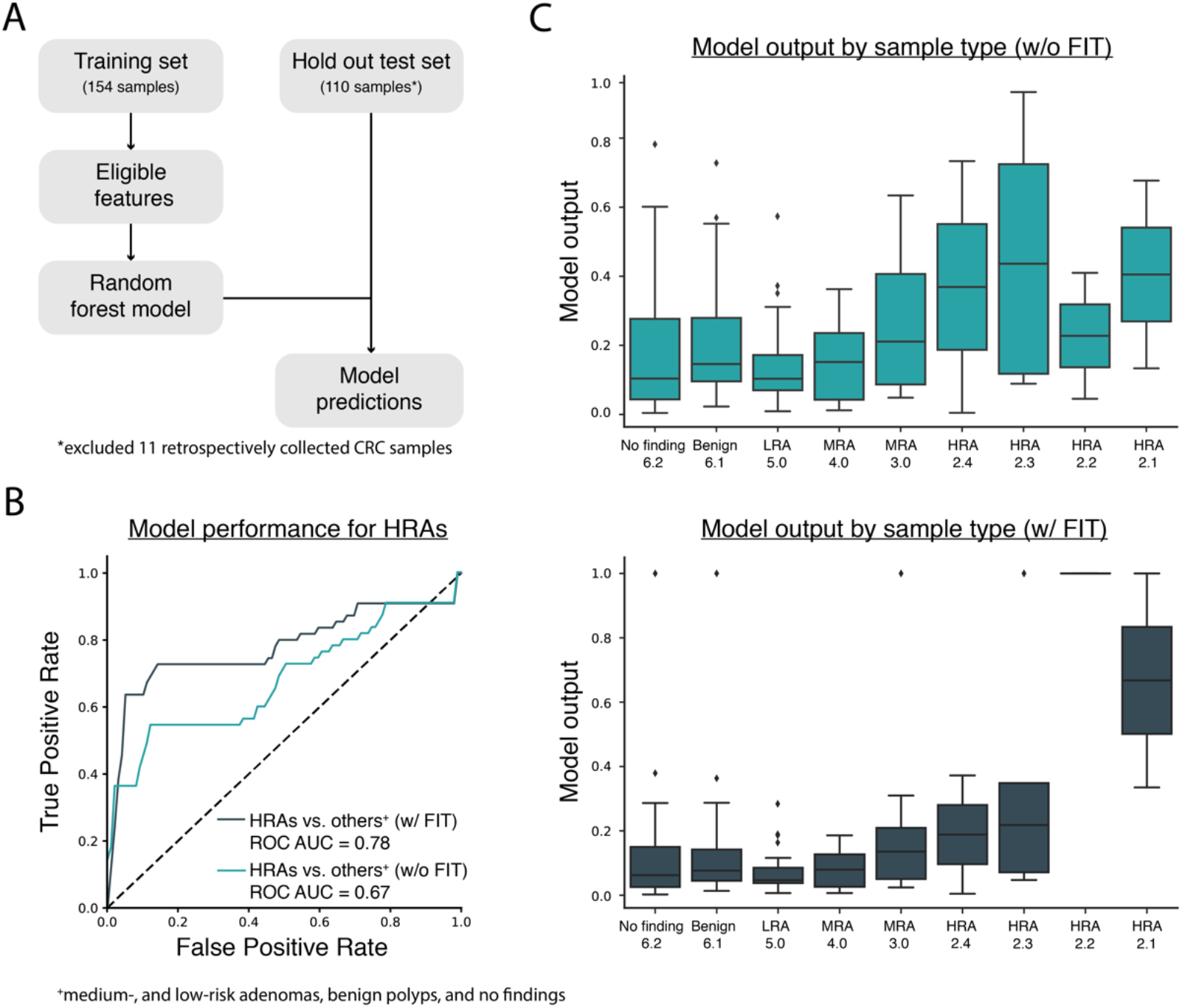
Model performance for the detection of HRAs based on the prospective hold out test set (n = 110 samples). **A)** A random forest model was created using the training set (n = 154 samples) and all 13 eligible features (see **Figure 2A**). The random forest model was employed on the prospective hold out test set (n = 110 samples) to determine model performance. Analysis was performed with and without using the FIT feature. When using the FIT, the model output was forced to 1 for FIT positive samples. **B)** The ROC curve shows model performance on the prospective hold out test set with and without the FIT feature. High-risk adenomas (HRAs) were considered positive and other findings (medium-risk adenomas, low-risk adenomas, benign polyps, no findings on a colonoscopy) were considered negative. **C)** Box plots show model output for each sample, parsed by sample type, for the prospective hold out test set. The top graph shows model predictions without the FIT feature and the bottom graph shows model prediction with the FIT feature. Sample type is ascending based on lesion severity (i.e., No finding 6.2 = least severe, HRA 2.1 = most severe) (see **Table 1**). The box plots represent quartiles, the bar represents the median value, the tails represent 95% of the data, and the dots indicate outliers (>2 standard deviations). *Abbreviations: FIT – fecal immunochemical test; ROC – receiver operator characteristic; AUC – area under the curve; LRA – low-risk adenoma; MRA – medium-risk adenoma; HRA – high-risk adenoma*

### Assessment of extent of model training using incremental downsampling analysis

The extent to which model performance was optimized was assessed using a downsampling analysis of the 154 samples in the training set (**see Methods**). The downsampling analysis showed a positive relationship between total number of samples used for training and performance on the hold out test set. When excluding FIT results, median ROC AUC for HRAs versus all other categories increased from 0.55 (30% of training data) to 0.67 (100% of training data) (**Supplementary Figure 4A**). When including FIT results, median ROC AUC for HRAs versus all other categories increased from 0.72 (30% of training data) to 0.78 (100% of training data) (**Supplementary Figure 4B**).

### Accuracy profile for CRCs and HRAs using 121-patient hold out test set

Predictions incorporating FIT results were also made for the 11 retrospectively collected stool samples from CRC patients. Using all 121 samples in this complete hold out test set, the model attained a ROC AUC of 0.94 for CRC versus all other categories (HRAs, MRAs, LRAs, benign polyps, and no findings on a colonoscopy) and a ROC AUC of 0.87 for CRC and HRAs versus all other categories (MRAs, LRAs, benign polyps, and no findings on a colonoscopy) (**Figure 4A**). When weighting cancer and HRAs to expected prevalence in a prospective screening population, the model attained a ROC AUC of 0.80 for CRC and HRAs versus all other categories (**Figure 4B**).

**Figure 4.**
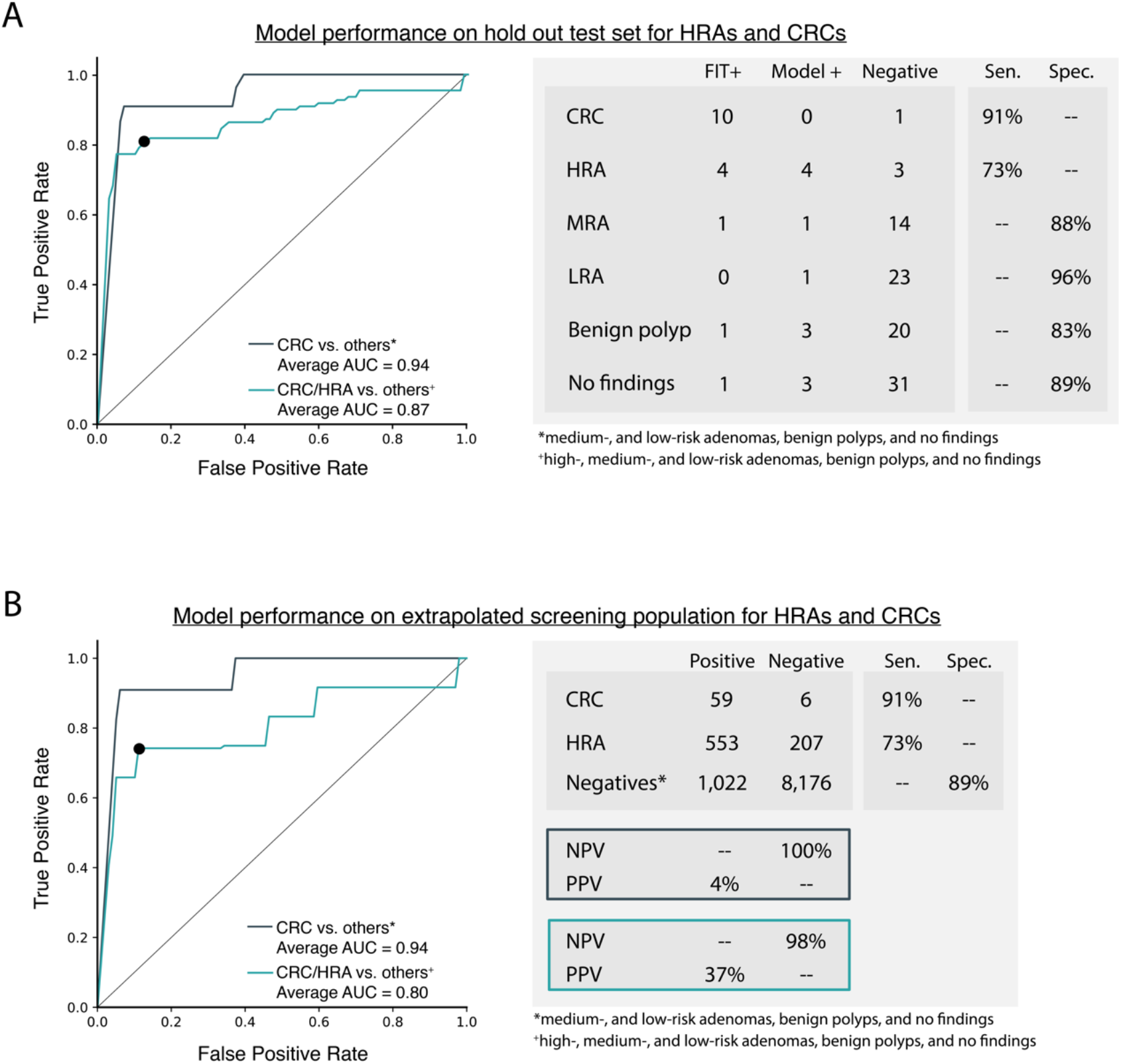
Model performance for the detection of CRC and HRAs based on the independent hold out test set (n = 121 samples) and extrapolation of model performance onto a screening population. **A)** The final model was employed on all samples in the hold out test set (including 11 retrospectively collected CRC samples). Model predictions could be positive based on the fecal immunochemical test (FIT+) or the random forest model prediction (Model+). Otherwise, the model prediction was considered negative (Negative). The sensitivity is shown for CRC and HRAs and specificity is shown for medium-risk adenomas, low-risk adenomas, benign polyps, and no findings on a colonoscopy. **B)** The model performance on the hold out test set was extrapolated to a generalized screening population to assess the negative predictive value and positive predictive value. *Abbreviations: FIT – fecal immunochemical test; ROC – receiver operator characteristic; AUC – area under the curve; Sen. – sensitivity; Spec. – specificity; LRA – low-risk adenoma; MRA – medium-risk adenoma; HRA – high-risk adenoma; CRC – colorectal cancer; NPV – negative predictive value; PPV – positive predictive value*.

Upon selection of the optimal point on the ROC curve to maximize sensitivity and specificity, the model demonstrated a 91% sensitivity for CRC (n = 11 samples) and a 73% sensitivity for HRAs (n = 11 samples) at an 89% specificity (n = 99 samples). Extrapolation of results onto a prospective screening population enabled calculation of the blended sensitivity for CRC and HRAs, the negative predictive value (NPV), and the positive predictive value (PPV). This extrapolated accuracy profile demonstrated a blended sensitivity for CRC and HRAs of 74%, a positive predictive value of 37%, and a negative predictive value of 98% (**Figure 4C**).

## Discussion

Currently, the most commonly used noninvasive screening alternative for CRC (Exact Sciences, Cologuard) offers a 92% sensitivity for CRC and a 42% sensitivity for HRAs, at an 87% specificity.^10^ Cologuard has received strong adoption, with 147,000 prescribing physicians and 934,000 tests completed in 2018. This test volume represents 4.1% of the total market for CRC screening, demonstrating the significant opportunity that exists for an improved noninvasive screening tool.^27^ Despite Cologuard’s strong commercial adoption, the test’s accuracy profile does not significantly improve upon existing, cheaper alternatives. One fecal immunochemical test (Polymedco, OC-Light S FIT) has a documented accuracy of 93% for CRC at a 91% specificity (estimated HRA accuracy is 25-35% at unknown specificity).^11^ Since Cologuard is a FIT-DNA test, it is clear that the addition of the nine stool-derived DNA biomarkers do not improve the CRC accuracy and offer insignificant improvement in HRA accuracy.

The FIT-RNA assay described here, attained a 91 % sensitivity for CRC, a 73% sensitivity for HRAs, and an 89% specificity for all other findings. This accuracy profile represents a significant improvement relative to all existing noninvasive alternatives. Specifically, the reported 73% sensitivity for HRAs represents a 74% improvement over Cologuard and the 74% blended sensitivity for CRC and HRAs represents a 61 % improvement over Cologuard. Improved sensitivity is attributable to the use of seRNA biomarkers, which offer several advantages relative to other stool- or blood-based biomarkers. First, seRNA biomarkers are derived from epithelial cells shed within the gastrointestinal tract. Therefore, the signal of the seRNA represents a homogenized sampling of the perilesional tissue, which is preferentially shed into the lumen and excreted in stool. Second, seRNA provides a concentrated and amplified signal that can be observed across multiple transcripts in a single pathway. This is a direct advantage over DNA-based biomarkers, which are limited to two copies per cell. Finally, the RNA transcriptome provides an assessment of the downstream molecular consequence of many precancerous variants regardless of tumorigenesis pathway. Therefore, a relatively small panel of seRNA biomarkers can be used to capture the vast genomic landscape that causes adenoma development. Leveraging seRNA biomarkers to attain high sensitivity for HRAs enables development of a diagnostic that would facilitate early intervention and prevention, rather than merely detection, of CRC.

Our use of a prospective study for biomarker selection and model development provides a true assessment of model performance in the desired screening population. This is a major advantage relative to the majority of research conducted in this space. For example, in 2012, Exact Sciences conducted a retrospective study with 678 patients (252 patients with CRC, 133 patients with HRAs, and 293 patients with no findings on a colonoscopy). Using only stool-derived DNA biomarkers and no FIT test, this study demonstrated an accuracy of 85% for CRC and an accuracy of 54% for HRAs at a 90% specificity.^28^ When comparing the impact of the DNA biomarkers in the prospective study relative to the retrospective study, it is clear that the retrospective study overestimated the DNA biomarkers’ sensitivity and specificity for both CRCs and HRAs. This overestimation is likely attributable to the exclusion of intermediate patient populations (i.e., MRAs, LRAs, and benign polyps) for marker selection and assessment. This problematic use of a retrospective cohort is also common in blood-based screening research. For example, cell-free DNA^29^ and circulating tumor cell^30^ studies cite high sensitivity and specificity on retrospectively selected samples from specific cohorts (CRC or HRAs, vs. no findings on colonoscopy). However, these studies have neither tested samples from intermediate patient populations nor tested samples from patients with confounding diseases or other cancers. The results described herein are not subject to aforementioned limitations. In this study, biomarker selection and accuracy profile development used samples from all patient populations within the CRC screening cohort. Specifically, patients with CRC or HRAs were considered positive and patients with all other findings (MRAs, LRAs, benign polyps, and patients with no findings on a colonoscopy) were considered negative. By overcoming common limitations of other development studies, we anticipate a consistent accuracy profile throughout development of the FIT-RNA test.

The features used for model development are unique relative to other biomarkers used for CRC screening. As expected, the 10 differentially expressed transcripts employed in the model have been previously implicated in CRC tumorigenesis. For example, *ACY1* is highly expressed in CRC tissue and is positively correlated with tumor stage.^31^ *SMAD4*^32^ and *CTNNB1*,^33^ are involved in the Wnt signaling pathway.^34^ Other selected transcripts are related to cell adhesion (i.e., *ACHE, DST*), cell growth (i.e., *EGLN2, ERBB2*), and cell death (i.e, *EDN1, TNFRSF10B*) (**Supplementary Table 1**). Novel features include the use of patient demographics and raw GAPDH values. It was observed that, relative to healthy patients, patients with MRAs, HRAs, and CRC had elevated raw *GAPDH* values. *GAPDH* is a commonly used epithelial housekeeping gene and it was hypothesized that increased total human RNA could be due to perilesional inflammation of the intestinal wall and subsequent sloughing of human cells into the lumen. Therefore, raw *GAPDH* values could provide a relative estimate of the total number of cells being sloughed and extracted during stool sample processing. Raw *GAPDH* was an effective biomarker in model development (Gini Importance score = 0.087) and assisted in detection of CRC and HRAs in the hold out test set. We also used age and smoking status for model development, which have known implications in tumor development but have never before been employed as features for CRC screening.

An unexpected observation was that the model output was correlated with disease severity in the hold out test set (**Figure 3C**). This correlation was a direct reflection of CRC biology and not trained as part of the model. Specifically, feature selection and model input used three categories (HRAs, MRAs, and all other), but directionality of categories (e.g., medium-risk adenomas are less severe than high-risk adenomas) and disease subsets (e.g., high-risk adenoma stratification based on adenoma size or level of dysplasia) were not available for model training. Given that the model output provided a well-scaled confidence in disease severity, it could be used to further stratify patient care. For example, if a patient’s model output is negative but is relatively high within the negative cohort, it might indicate that the patient has an MRA that would require a higher frequency of screening relative to a negative patient with a lower score.

The biomarker panel for CRC and HRA detection used only 11 seRNA biomarkers, 1 FIT, and 2 demographic features; therefore, supplementing the existing panel with additional biomarkers could further improve the accuracy profile. This could be accomplished by adding additional seRNA expression markers or by using expressed somatic variants for known transcripts (e.g., *APC, KRAS, or TP53*). Use of a FIT, transcript expression, and expressed variants could provide a comprehensive panel that encapsulates a variety of biomarkers in the same assay. Supplementing the existing panel with new biomarkers could also improve medium-risk adenoma (MRA) detection. In this study, we observed that MRAs exhibit a unique gene expression pattern that could be included in the model for sensitive detection of MRAs. Based on preliminary research presented here, we believe that seRNA biomarkers could be used to simultaneously detect CRC, HRAs, and MRAs, which could provide population health benefits in a prospective screening population.

The low incidence of CRC in a screening population made it difficult to prospectively obtain stool samples from patients with CRC prior to a colonoscopy. To understand model performance on CRC samples, it was required to supplement the prospective collection with 11 samples that were obtained retrospectively from patients who had been diagnosed with CRC but who had not yet undergone surgical resection or treatment. It is possible that model results from this cohort would be different if the samples were obtained prior to screening colonoscopy and biopsy. That being said, the CRC accuracy profile reported here directly reflects the known accuracy profile of the OC-Light S FIT (Polymedco), which was the FIT used in this study. Therefore, we expect that the reported CRC accuracy profile (91% sensitivity) would be comparable in a prospective study.

Given that most CRC samples were collected retrospectively, the hold out test set was enriched for CRC. Measuring the negative predictive value (NPV) and positive predictive value (PPV) required extrapolating sensitivity and specificity results to a generalized screening population. This extrapolation analysis still showed that a model with the aforementioned accuracy profile would be a robust method for HRA and CRC detection with high blended sensitivity (74%) and high NPV (98%).

Limitations of this study include use of a single organization for sample collection (3 endoscopy sites), use of a hold out test set obtained from the same collection sites, and a limited number of CRCs and HRAs in the hold out test set (n = 22 samples). This study is also limited by use of a retrospective cohort for CRC detection as described above. Future studies will increase breadth and diversity of collection sites and will increase total number of samples with positive findings in the hold out test set.

In summary, this research validates the use of seRNA biomarkers for sensitive detection of CRC and HRAs. We show that these biomarkers can provides accurate partitioning of CRCs and HRAs from all other negative findings. Our novel use of a prospective cohort for biomarker selection and model development creates the first pre-FDA approval study with an accuracy profile that can be reproduced as study size increases. Use of a prospective screening cohort also provides insight on detection of smaller pre-malignant lesions (i.e., MRAs), detection of which has been challenged by existing noninvasive screening options. These data provide evidence that seRNA biomarkers could significantly impact the ability to noninvasively screen for colorectal neoplasms for the millions of Americans that are noncompliant with existing screening guidelines.

## Acknowledgements

We would like to thank our scientific advisors: Dave Messina, Ira Kodner, and Phil Needleman. This research would not have been possible without support and assistance from Accelerate St. Louis, Arch Grants, BioSTL, BioGenerator Labs, Sling Health St. Louis, The Global Impact Award, Cofactor Genomics, The Wharton School, and Washington University School of Medicine.

## Author Contributions

EB and YK contributed to the study concept and design; EB, YK, AB, KC, and EW contributed to the acquisition, analysis, and interpretation of data; EB, YK, AB, KC, KK, and EW contributed to drafting of the manuscript; MG, AC, and OG contributed to critical revision of the manuscript; EB, YK, AB, and KC contributed to statistical analysis; EB, YK, and AB obtained funding; MG, AC, and OG contributed to study supervision. All authors have seen and approved the final draft.

## Financial Contributions

This research was funded by Geneoscopy LLC (St. Louis, MO), which is a startup company developed from technology at the Washington University School of Medicine and grants from Accelerate St. Louis, Arch Grants, BioGenerator Labs, The Global Impact Award, and The Wharton School. Malachi Griffith is supported by the NHGRI under award number K99HG007940. Aadel Chaudhuri is supported by the Washington University SPORE in Pancreatic Cancer Career Enhancement Program and the Cancer Research Foundation Young Investigator Award. Obi Griffith is supported by the NCI under award number K22CA188163. The content of this manuscript is solely the responsibility of the authors and does not necessarily represent the official views of the NIH.

## Abbreviations

AUC: area under the curve ANOVA analysis of variance
CRC: colorectal cancer
DE: differentially expressed
FC: fold-change
FIT: fecal immunochemical test
HRA: high-risk adenoma
LRA: low-risk adenoma
MRA: medium-risk adenoma
NPV: negative predictive value
PPV: positive predictive value
ROC: receiver operating characteristic
seRNA: stool-derived eukaryotic RNA

