## Supplementary Tables and Figures for "Stool-derived eukaryotic RNA biomarkers for detection of high-risk adenomas"

##### Supplementary Table 1. Features employed in model development

| Feature | Feature Type | Description | Gini Importance |
|---|---|---|---|
| ACY1 | Normalized Expression | ACY1 catalyzes hydrolysis of acylated proteins into L-amino acids and acyl groups and regulates cancer cell proliferation. ACY1 mRNA is expressed in CRC tumor tissue and expression is positively correlated with tumor stage. | 0.134 |
| TNFRSF 10B | Normalized Expression | TNFRSF10B is a cell surface receptor that mediates the extrinsic pathway of apoptosis. Expression has been implicated in cancer treatment. Specifically, exposure to therapy results in endoplasmic reticulum stress and apoptosis. | 0.130 |
| DST | Normalized Expression | DST is a cytoskeletal linker protein essential for maintaining the cytoskeletal integrity of neurons. Elevated expression has been demonstrated in melanomas. | 0.091 |
| GAPDH | Raw Expression | GAPDH plays a role in glycolysis and is a commonly used as a housekeeping gene. It has been shown to be overexpressed in tumors of some cancers. | 0.087 |
| EDN1 | Normalized Expression | EDN1 governs cell proliferation, apoptosis, migration, and chemo-resistance. Expression contributes to tumorigenesis through autocrine and paracrine signaling. Increased levels of EDN1 have been detected in CRC patients. | 0.079 |
| EGLN2 | Normalized Expression | EGLN2 is involved in oxygen homeostasis. Indels in the proximal promotor have been associated with significantly increased risk for CRC. | 0.074 |
| Age | Demographic | Age has been demonstrated to be directly correlated with CRC development. | 0.073 |
| ACHE | Normalized Expression | ACHE terminates signal transduction at the neuromuscular junction via hydrolysis of acetylcholine. Low expression is associated with aggressiveness and recurrence in hepatocellular carcinoma. | 0.068 |
| Smoking Status | Demographic | Smoking has been demonstrated to be directly correlated with CRC development. | 0.060 |
| ERBB2 | Normalized Expression | ERBB2 is a member of the epidermal growth factor family. It is involved in the transcription of rRNA to enhance protein synthesis and cell growth. Overexpression contributes to tumorigenesis with previous implication in CRC. | 0.055 |
| SMAD4 | Normalized Expression | SMAD4 plays a role in the TGF-B signaling pathway as a transcriptional regulator. Loss of SMAD4 leads to tumor growth and progression, it is therefore an important marker for pancreatic and other cancers. | 0.054 |
| CTNNB1 | Normalized Expression | CTNNB1 plays a role in cell adhesion, intercellular communication, cell proliferation and cell differentiation. Its action is mediated through the Wnt signaling pathway. Somatic mutations result in excess beta-catenin buildup and tumor development. These mutations are often associated with CRCs. | 0.050 |
| PER3 | Normalized Expression | PER3 is involved in regulation of the circadian cycle; plays an important role in sleep-wake timing, and sleep homeostasis. Decreased expression associated with increased susceptibility to cancer development. | 0.045 |

### Supplementary Figures

**Supplementary Figure 1. The technical sequencing experiment used the same stool sample, extraction, and library preparation. Library preparations were aliquoted into two wells and sequenced on two different NextSeq runs.**

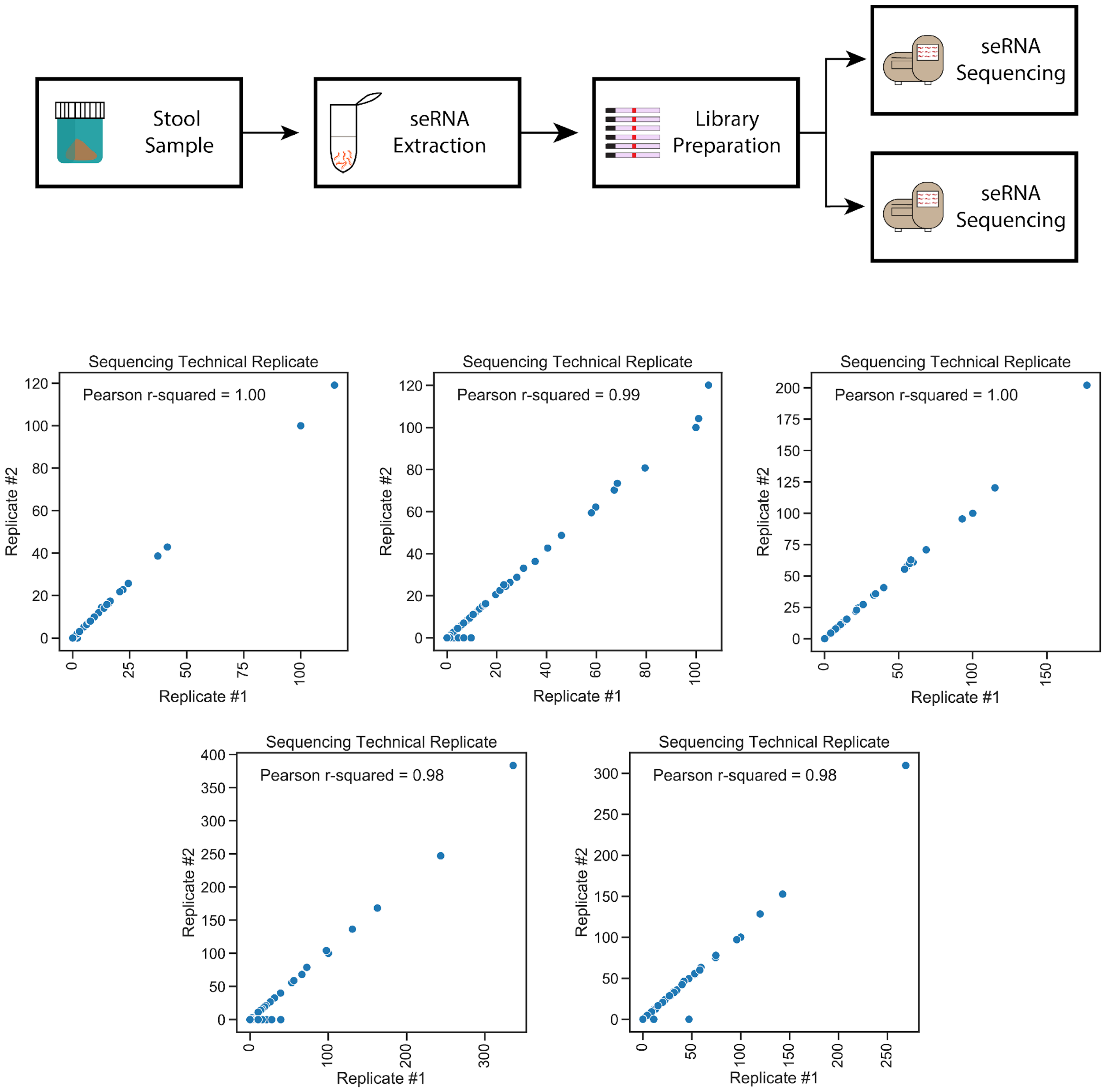

**Supplementary Figure 2. The biological library preparation experiment assessed different stool samples extracted at different times, prepared using different library preparation parameters (200 ng / 30 cycles vs. 400 ng / 28 cycles) but sequenced on the same NextSeq run.**

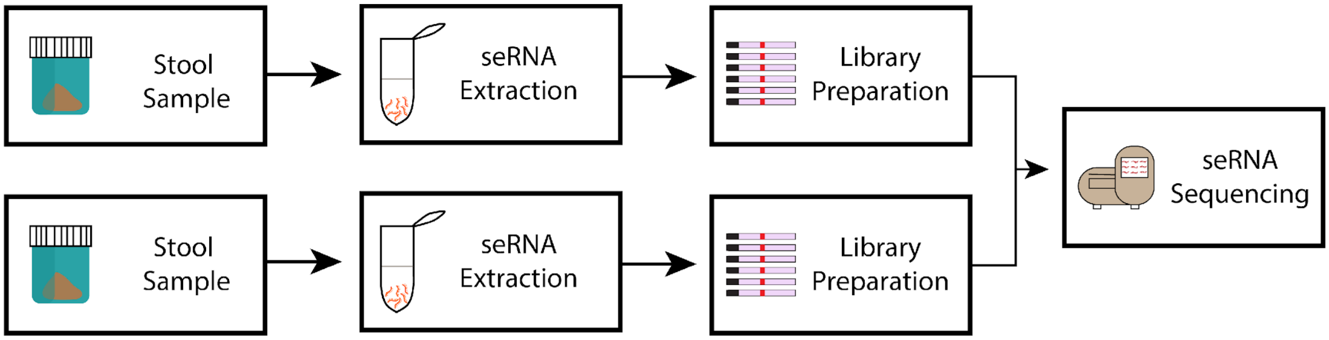


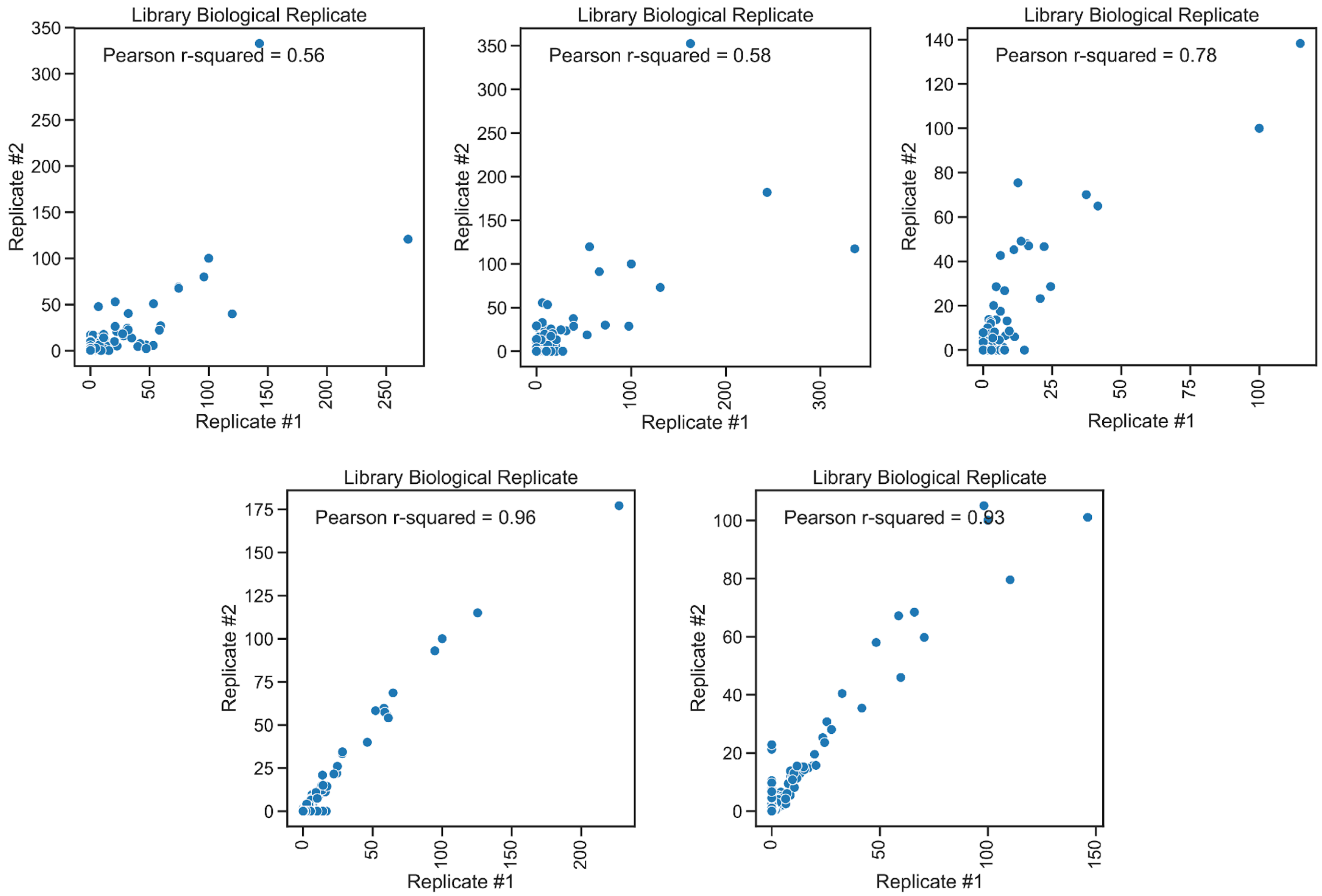

**Supplementary Figure 3. The biological sequencing experiment assessed different stool samples extracted at different times, prepared with the same library preparation parameters (400 ng / 28 cycles), and sequenced on different NextSeq runs.**

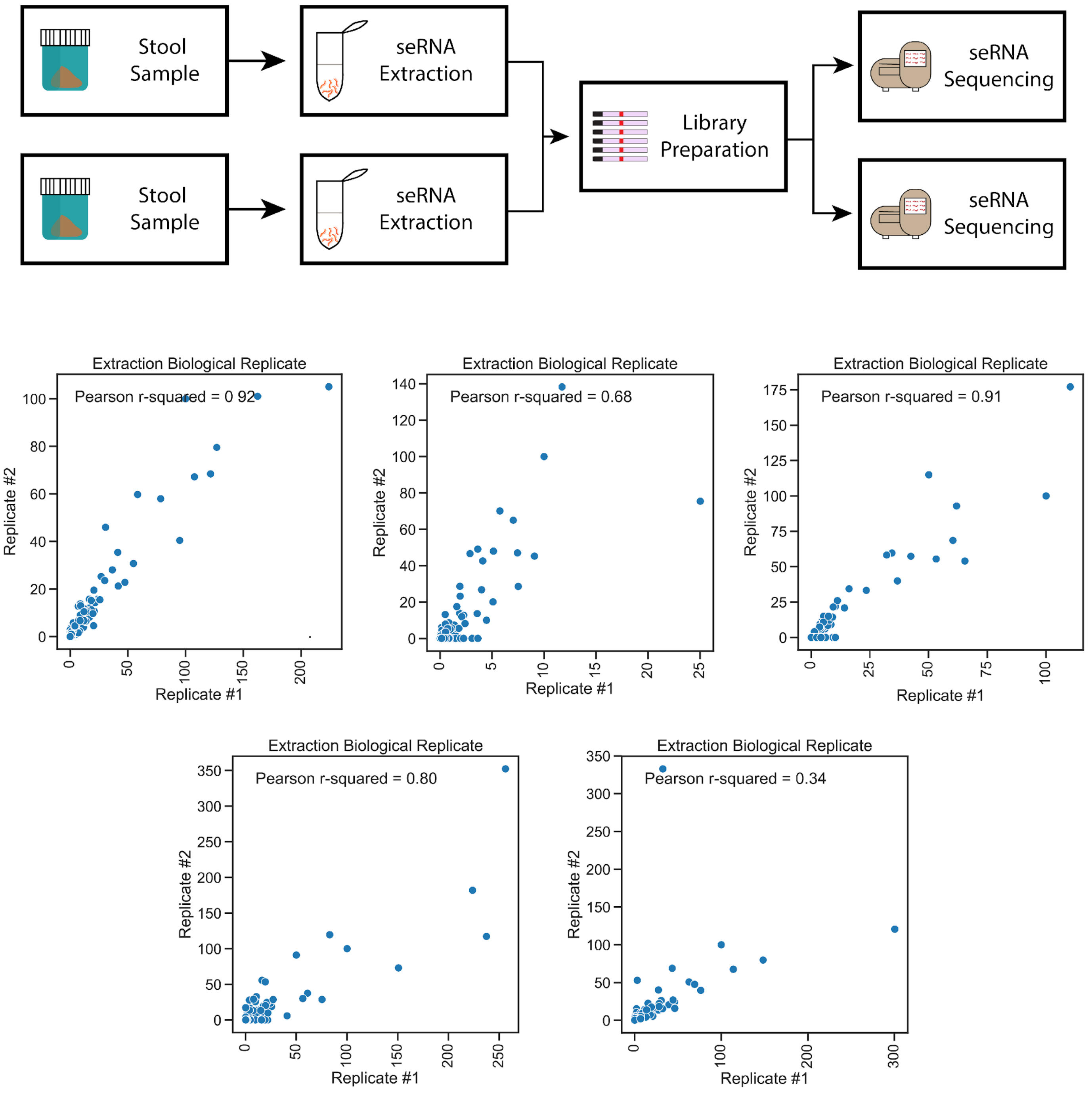

**Supplementary Figure 4. Incremental downsampling analysis performed on hold out test set with and without FIT results**. Box plots show model performance for each downsampled fraction. The box plots represent quartiles, the bar represents the median value, the tails represent 95% of the data, and the dots indicate outliers (>2 standard deviations). Models were built using downsampled versions of the training set (n = 154 samples) and were employed on the prospective hold out test set (n = 110 samples). For each increment, this downsampling and testing process was run ten separate times. The model performance as measured by the ROC AUC for all runs at each increment was plotted. **A)** Incremental model performance on the hold out test without the FIT feature. **B)** Incremental model performance on the hold out test with the FIT feature.

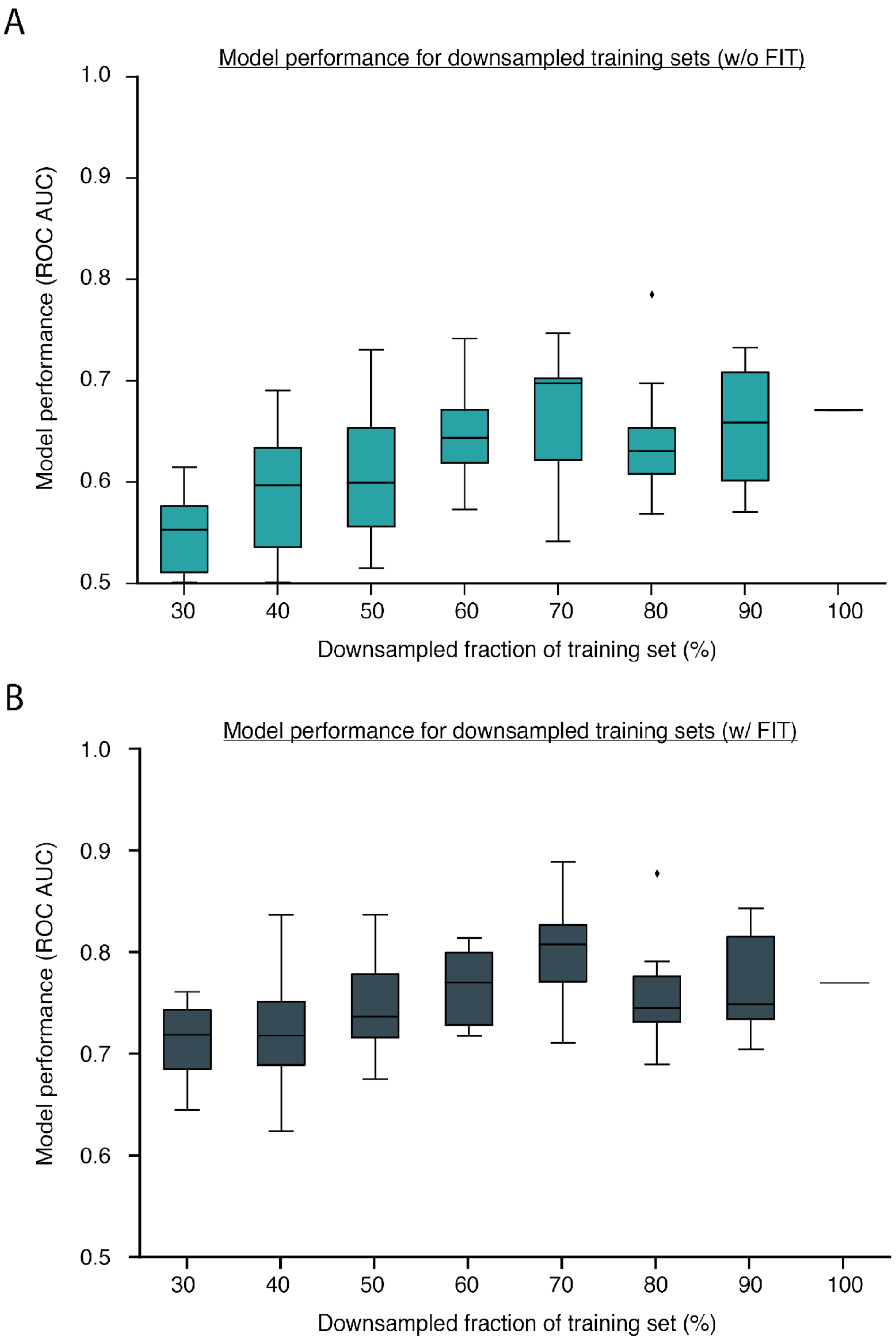
