## Supplementary Methods for "Stool-derived eukaryotic RNA biomarkers for detection of high-risk adenomas"

*Eligible patients and sample collection*

Stool samples were obtained by the Digestive Diseases Research Core Center (DDRCC) at WUSM. All patients were sent a stool sample collection kit by mail and returned the kit via courier to the DDRCC. Clinical data (e.g., demographic information, colonoscopy results, etc.) were collected by the DDRCC. Each sample was tested for blood in the stool using a commercially available FIT (Polymedco, OC-Light S FIT)[1] prior to being frozen at -80°C.[2] Each patient recruited for the study had a colonoscopy performed and those with positive findings underwent biopsy and subsequent histopathologic review to determine neoplastic classification. Adenoma classification was stratified based on histopathology (benign vs. pre-malignant), number of polyps, size of polyps, and differentiation. Cancer classification was stratified based on the American Joint Committee on Cancer (AJCC) 7 TNM system.[3] If the patient had no findings during the colonoscopy, he or she was labeled as healthy.

*Development of a training set and a testing set*

In total, 275 stool samples were collected. 154 prospectively collected stool samples were used as a training set and 110 prospectively collected stool samples were used as a hold out test set. 11 retrospectively collected stool samples from CRC patients were also included in the hold out test set. The training set and hold out test set were evaluated for categorical, demographic, and handling differences using a t-test (population means) or z-test (population frequencies) and significance was indicated if the p-value was less than 0.05.

*Stool sample enrichment for human RNA from exfoliated enterocytes using differential centrifugation*

Stool samples were aliquoted into 50mL conical tubes and filled to 45mL using suspension buffer (10mM Tris, 1mM EDTA, 0.005% Tween-20, 80U RNase Inhibitor, pH 7.5). Samples were homogenized and subjected to differential centrifugation using a swing-bucket centrifuge for 10 minutes at 4°C to separate the homogenate into a human cell layer below an enriched bacterial supernatant. The supernatant was discarded to eliminate bacterial noise and the pellet was suspended into a guanidine thiocyanate buffer to lyse the enterocytes and expose the human biomarkers.[4] From the lysate layer containing enriched human nucleic acids, 2mL of the solution was purified using an automated NucliSENS easyMag (bioMérieux, Durham, NC).[5,6] The final solution was subjected to Baseline-ZERO DNase treatment (Epicentre Technologies, Madison, WI) and clean-up using the automated NucliSENS easyMag.[7]

*Custom amplicon panel development*

The custom amplicon panel was developed using previously conducted research on 349 samples and literature review. First, transcripts were selected based on a microarray experiment (Accession #GSE99573).[8] For this experiment, total seRNA was extracted from stool samples and expression was assessed using Affymetrix Human Transcriptome Array 2.0 (ThermoFisher Scientific, Waltham, MA). Microarray expression profiles derived from patients with CRC or pre-malignant adenomas (diseased cohort) were compared to expression profiles from patients with no findings on a colonoscopy (healthy cohort). Transcripts with significant differential expression (p<0.03) were selected for the amplicon panel. Additional transcripts were selected based on a NanoString experiment.[8] Again stool samples were obtained from a diseased cohort and a healthy cohort. Total seRNA was extracted from stool samples and expression was assessed using the nCounter® PanCancer Pathways Panel (NanoString, Seattle, WA) and the nCounter® PanCancer Progression Panel (NanoString, Seattle, WA). Differentially expressed transcripts were identified by comparing the diseased cohort to the healthy cohort using the nSolver differential expression analysis platform. Finally, the literature was evaluated for additional transcripts implicated in CRC. This included searching ClinVar[9], COSMIC[10], CIViC[11], and other pertinent studies.[12,13] A custom amplicon panel was developed for targeted enrichment using the Illumina DesignStudio (**Figure 2A**).

*seRNA quality check and sequencing*

seRNA integrity was determined using the Agilent 2100 Bioanalyzer (Agilent Technologies, Santa Clara, CA) and mass was determined using a Qubit Fluorometer (ThermoFisher, Waltham, MA). Samples required >200ng of RNA to be eligible for library preparation. Libraries were prepared using a TruSeq Targeted RNA Custom Panel (Illumina, San Diego, CA). Sequencing was performed using the NextSeq 550 System (Illumina, San Diego, CA). A PhiX spike-in was used for quality control.

*Samples selected for technical and biological replicates*

Five patients were used to assess reproducibility of seRNA extraction and sequencing. Across these patients, three separate reproducibility experiments were performed: technical sequencing replicate, biological library preparation replicate, and biological sequencing replicate. Pearson r correlation was used to assess change in transcript expression for all replicates across all amplicons in the custom panel.

The technical sequencing replicate assessed reproducibility of independent sequencing runs for transcript expression quantification. For this experiment, a single sample was subjected to extraction and library preparation. Subsequently, two separate aliquots of the library preparation were sequenced on different sequencing runs.

The biological library preparation replicate experiment assessed reproducibility of two different library preparations for transcript expression quantification. For this experiment, a stool sample from the same bowel movement was split into 2 aliquots that were homogenized and extracted separately. One sample used 400 ng of library input and was subjected to 28 cycles of polymerase chain reaction (PCR) amplification. The second sample used 200 ng of library input and was subjected to 30 cycles of PCR amplification. Both replicates underwent parallelized sequencing on the same sequencing run.

The biological sequencing replicate experiment assessed reproducibility of independent sequencing runs for transcript expression quantification. For this experiment, a stool sample from the same bowel movement was split into 2 aliquots that were homogenized and extracted separately. Both samples used 400 ng of library input and were subjected to 30 cycles of PCR amplification. The samples were subsequently sequenced on independent sequencing runs.

*Quantification of seRNA expression*

Raw sequencing reads were aligned to amplicons of interest using the reference genome (GRCh38) via HISAT2.[14] Samples required >100,000 aligned reads for study eligibility. For each sample, raw amplicon expression was normalized to *GAPDH*, the internal housekeeping gene, such that reported expression equates to amplicon read count per million mapped *GAPDH* reads. Raw *GAPDH* values were used as a measure for total eukaryotic RNA in each sample. Raw *GAPDH* values were eligible as a feature for model development.
